# Bioaugmentation strategies based on bacterial and methanogenic cultures to relieve stress in anaerobic digestion of protein-rich substrates

**DOI:** 10.1101/2023.12.11.571062

**Authors:** Sara Agostini, Luca Bucci, Davide Doni, Paola Costantini, Ameya Gupte, Bettina Müller, Fabrizio Sibilla, Marina Basaglia, Sergio Casella, Panagiotis G. Kougias, Stefano Campanaro, Lorenzo Favaro, Laura Treu

## Abstract

Anaerobic co-digestion of protein-rich substrates is a prominent strategy for converting valuable feedstocks into methane, but it releases ammonia, which can inhibit methanogenesis. This study developed a cutting-edge combined culturomic and metagenomic approach to investigate the microbial composition of an ammonia-tolerant biogas plant. Newly-isolated microorganisms were used for bioaugmentation of stressed batch reactors fed with casein, maize silage and their combination. A co-culture enriched with proteolytic bacteria was isolated, selected and compared with the proteolytic collection strain *Pseudomonas lundensis* DSM6252. The co-culture and *P. lundensis* were combined with the ammonia-resistant archaeon *Methanoculleus bourgensis* MS2 to boost process stability. A microbial population pre-adapted to casein was also tested for evaluating the digestion of protein-rich feedstock. The promising results suggest combining proteolytic bacteria and *M. bourgensis* could exploit microbial co-cultures to improve anaerobic digestion stability and ensure stable productivity even under the harshest of ammonia conditions.

**Graphical abstract:** 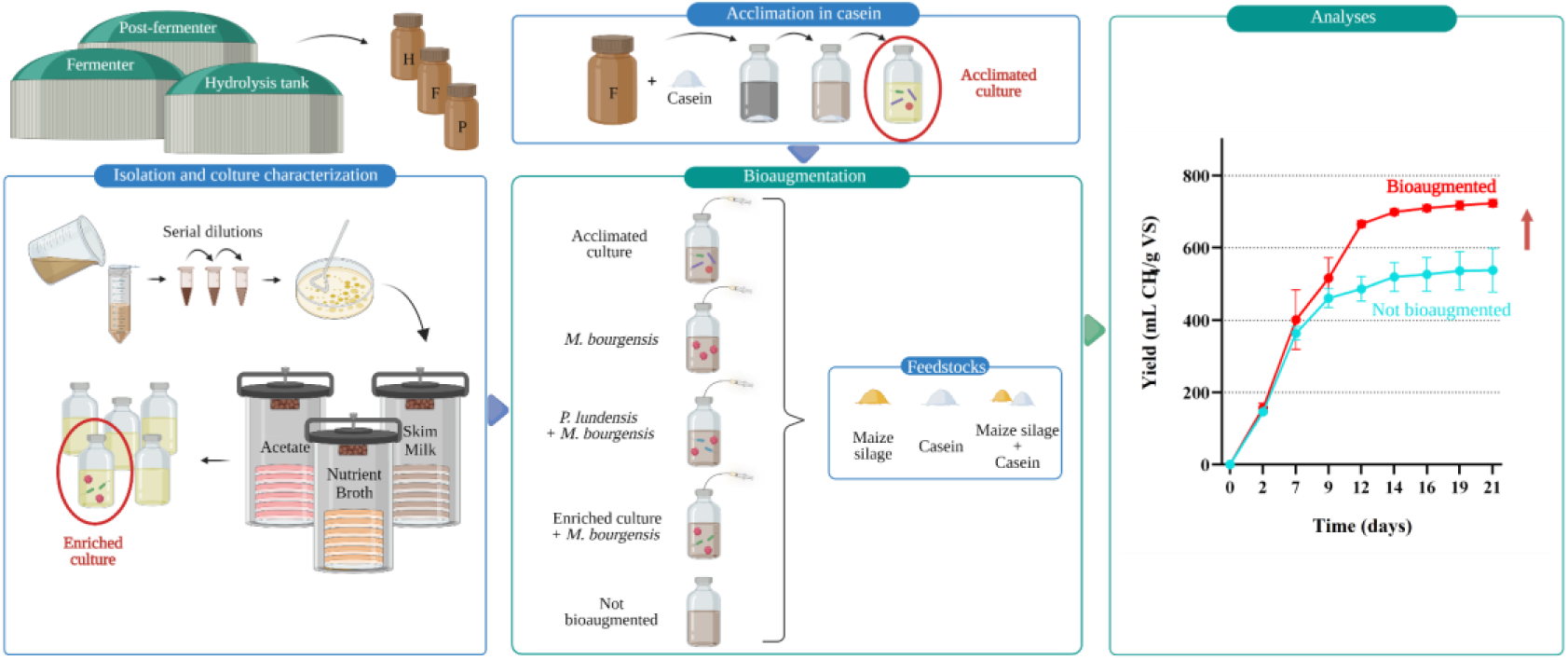

**Highlights:**

- High ammonia release from protein-rich substrates inhibits anaerobic digestion
- Newly isolated bacterial strains from anaerobic digester were obtained
- Proper bioaugmentation alleviates stress in casein and maize silage co-digestion
- Bioaugmentation with a hydrolytic/hydrogenotrophic co-culture enhances CH_4_ yields

## 1. Introduction

Renewable energy sources capable of replacing fossil fuels are considered fundamental strategies to face the energy and environmental crisis (Li et al., 2020). Biogas produced by digesting organic material is a fuel gas rich in methane, rated as one of the unquestioned protagonists of the green economy (Liu et al., 2022). It can generate heat and energy (Scarlat et al., 2018) and, when purified through biogas upgrading, can be used as a vehicle fuel or injected into the gas grid (Prussi et al., 2019). The anaerobic digestion (AD) process, aside from generating biogas and fertilising digestate, is an interesting solution for the management and recycling of waste materials and residues from agriculture and industry (e.g. manure and crop biomass), municipal solid waste (OFMSW), sewage sludge, and urban wastewater (Achinas et al., 2017; Brojanigo et al., 2022).

AD is a biologically mediated process in which the whole microbial community cooperates in the degradation of organic substrates, resulting in biogas production. It consists of a series of interconnected metabolic reactions grouped in four main activities: hydrolysis, acidogenesis, acetogenesis and methanogenesis (Venkiteshwaran et al., 2015). The latter is carried out only by archaea through three main pathways: hydrogenotrophic, acetoclastic and methylotrophic. Hydrolysis and acidogenesis are characterised by the abundance of different species potentially competing in metabolising the substrate. On the contrary, in the last two passages there is a lower functional redundancy level (Treu et al., 2018), where archaea and acetogenic bacteria tend to cooperate, by establishing syntrophic relationships.

The functionality of a biogas system is strictly related to the microbial community (Wu et al., 2022). The knowledge of microbial hierarchy, particularly in AD systems, has been recently obtained with high throughput sequencing techniques but some limitations towards the full microbiome picture are still present (Campanaro et al., 2018). Microbial species present at low concentrations are difficult to be detected, a threshold value for the taxonomic assignment at the species level is difficult to be set (Sankar et al., 2015), and some copies of the 16S rRNA gene can be acquired by horizontal gene transfer (Zhi et al., 2012). For these reasons, culturomics is considered a valid and complementary strategy to metagenomics for microbial studies. However, culturable microbes represent only a small fraction of the microbiota, a condition known as “the great plate count anomaly” (Amann et al., 1995). Many estimates were proposed in the past years to quantify the unculturable majority, and, depending on the environment, approximately 95–99.9% of the organisms of interest is estimated to not be readily culturable (Lelie et al., 2012).

In numerous organic wastes, such as those generated by whey, cheese, fish, and some vegetables industries, proteins are abundantly present (Sasaki et al., 2011; Treu et al., 2019). During AD these macromolecules are converted into ammonia, carbon dioxide and hydrogen (Ling et al., 2021). High ammonia concentrations in the system are estimated to be the primary cause of process inhibition (Chen et al., 2008). Although NH_3_ concentrations below 0.2 g/L are necessary for the microbial cells to grow, levels higher than 2 g/L can hinder biogas production (Yang et al., 2018). In particular, ammonia concentrations ranging from 3.5 to 7 g/L were reported to drastically reduce methanogens activity (Ling et al., 2021).

The inhibiting effects of ammonium are related to its hydrophobic form which penetrates the membranes causing intracellular damage (Ling et al., 2021). For this reason, FAN (Free Ammonia Nitrogen: NH_3_) is more toxic than TAN (Total Ammonia Nitrogen: NH_3_ + NH_4_^+^). Since the two forms are in dynamic equilibrium, their concentrations depend on pH and temperature (Cai et al., 2021): the increase of both these parameters leads to higher levels of free ammonia. Indeed, thermophilic cultures have been shown to be more sensitive to ammonium stress (Sung and Liu, 2003). Within the microbial population, methanogens are the most sensitive regarding this stress, as their metabolic activity is reduced and consequently, fermentation intermediates (H_2_ and volatile fatty acids) accumulate in the system leading to process inhibition (Shi et al., 2016). In particular, acetoclastic archaea are those having the highest sensitivity to ammonia (Bi et al., 2021). When acetoclastic methanogenesis can no longer be performed, a shift in the acetate utilisation occurs: the oxidation of acetate to H_2_ and CO_2_ by syntrophic acetate-oxidising bacteria (SAOB, more robust to ammonia stress) rises, sustaining hydrogenotrophic methanogenesis (Schnürer and Nordberg, 2008; Yenigün and Demirel, 2013).

To solve the problem of ammonia inhibition in AD, different solutions have been tested, including for example, the control of process parameters, co-digestion to adjust C:N ratio, microbial acclimatation and the addition of adsorbent materials (Palù et al., 2022; Tian et al., 2018b; Yan et al., 2021). Some of these approaches, such as pH control and temperature regulation, proved to be inefficient or economically expensive. On the contrary, the adaptation of the methanogenic population to high levels of ammonium, and the correlated change in the methane production pathways, was found to be effective (Tian et al., 2018b). An interesting solution can be represented by the bioaugmentation, thus enriching the AD system with a microbial consortium or a pure culture preadapted and/or tolerant to ammonia-related stress (Fotidis et al., 2014; Yan et al., 2020; Lovato et al., 2020).

Towards the efficient AD of protein-rich substrates, this study aimed to isolate novel bacterial strains or co-cultures to be efficiently adopted when the ammonia released from protein hydrolysis reaches microbes-inhibiting levels. An interesting co-culture with a proteolytic bacterium was characterised and used to facilitate the digestion of casein in AD settings. Several bioaugmentation strategies were then pursued exploiting the novel and promising co-culture as well as strains of a proteolytic bacterium or a hydrogenotrophic archaea obtained from cultures collection, in addition to a culture acclimated to casein. Variations in methane yields and environmental parameters, such as pH and ammonium levels, were then monitored to assess the bioaugmentation efficiency in a protein-fed AD process.

## 2. Materials and methods

### 2.1 Sampling of an industrial-scale biogas plant

The mesophilic (42°C) biogas plant considered in this study is located in Chiari (Brescia, Italy), and equipped with three digesters: hydrolysis (H), fermenter (F) and post-fermenter (P). At the sampling time the plant was fed with a mixture of chicken manure, maize silage and pig slurry. The substrate mix is transferred from the pre-hydrolysis tank to the first fermenter, where the main digestion takes place, and subsequently to the post-fermenter where the residual materials are degraded. The biogas plant has been working for years in stable conditions with an average ammonium level of 5.5 g NH_4_^+^-N/L, a pH of 8.0 and a Flüchtige Organische Säuren/Totales Anorganisches Carbonat (FOS/TAC) ratio of 0.23. Samples from each digester were kept at 42°C to perform species isolation, and one aliquot for each of them was sequenced for molecular characterisation of the microbiota.

### 2.2 DNA extraction and 16S rRNA sequencing

Genomic DNA of the microbial communities collected from H, F and P digesters, together with pure microbial strains, was extracted with the DNeasy PowerSoil kit (QIAGEN, Germany). In order to recover the pellet from microbial isolates, samples of 10 mL were previously centrifuged at 4000 rpm, 4°C for 15 min. The concentration and quality of purified genomic DNA were estimated using NanoDrop 2000 (ThermoFisher Scientific, MA, USA) and Qubit 2.0 fluorometer (Invitrogen, MA, USA), using the Qubit dsDNA High Sensitivity assay kit (Invitrogen).

Amplification of hypervariable region V4 of the 16S rRNA gene of digesters samples was performed using the degenerated primers pair 515F (5′-GTGYCAGCMGCCGCGGTAA-3′) and 806R (5′-GGACTACNVGGGTWTCTAAT-3′) as suggested by Parada et al., 2016.

For isolates identification, the combination of universal primer 10F (5′-AGTTTGATCCTGGCTCAG-3′) (Youssef et al., 2015) and degenerated primer 806R were used to amplify the rRNA 16S gene sequences. The PCR conditions used for all the reactions were: 32 cycles of a program with a denaturation phase of 45 sec at 98°C, an annealing of 40 sec at 58°C, and an elongation of 50 sec at 72°C. A final extension phase at 72°C for 5 min was performed at the end of the amplification.

Processing and sequencing of the H, F and P amplicons were performed with the Illumina MiSeq paired-end platform at the NGS facility of the Biology Department of the University of Padova (Padova, Italy). Raw reads (250+250 bp) have been submitted to the sequence read archive database (SRA) of NCBI (https://www.ncbi.nlm.nih.gov/sra) under the BioProject PRJNA866107 with accession numbers SRX16984443, SRX16984444, and SRX16984445. The amplicons of the 10-806 region of the 16S rRNA gene of isolates were sequenced with the Sanger method at BMR genomics S.r.l. (Padua, Italy), after being purified using AMPure XP beads (Beckman Coulter, CA, USA) to remove the primers following the instructions of the supplier.

### 2.3 Bioinformatic analyses

Raw reads in fastq format from H, F and P were analysed with CLC Genomics workbench software V.20.0.4 with microbial genomics module plug-in (CLC Bio, QIAGEN), using SILVA 138 as the amplicon reference database. The taxonomic assignment of each Operational Taxonomic Unit (OTU) and Sanger sequence was manually verified with similarity search through an alignment with NCBI Nucleotide Blast (https://blast.ncbi.nlm.nih.gov/Blast.cgi), using 16S ribosomal RNA sequences (Bacteria and Archaea) as database. Multi experiment viewer software (MeV 4.9.0) (Saeed et al., 2003) was used for visual representation of the high abundant OTUs as a heat map.

The Sanger sequences were visualised with the Chromas 2.6.6 software (http://www.technelysium.com.au/chromas.html).

### 2.4 Isolation and cultivation of microorganisms

Enrichment and isolation of the microorganisms were carried out under an anaerobic glove box with N_2_ atmosphere (MBRAUN MB200B, Germany). To isolate the highest number of bacterial strains, basal anaerobic (BA) medium (Angelidaki et al., 1990) was specifically supplemented with three different carbon sources: nutrient broth (NB) powder (No. 3 SigmaAldrich, MO, USA, 13 g/L) to support the growth of general anaerobic bacteria, acetate (2.5 g/L) with the aim of isolating archaea and/or SAOB, and skim milk (SM) powder (SigmaAldrich, 10 g/L) to facilitate the selection of proteolytic microbes. Yeast extract (YE) (LP0021, 26 Oxoid Limited, Cheshire, England) and a double volume of vitamin solution, with respect to that suggested in the BA medium preparation protocol, were added to better sustain the microbial growth.

The three samples from H, F and P reactors were filtered to remove larger particles. The suspensions were then serially diluted, plated on agar medium with acetate, NB or SM as carbon source and incubated at 42°C under anaerobic conditions for 21 days, after which CFU (colony forming units) were determined. Anaerobic conditions were obtained in anaerobic jars (Oxoid) flushed with CO_2_ and H_2_ gas in a ratio of 1:4. All steps were conducted in triplicates.

From each sample, colonies were purified through consecutive streakings. Colonies of interest were then inoculated in liquid medium with the same carbon source used for their isolation, and the volumes were subsequently increased to the final working volume of 50 mL in 120 mL bottles. Bottles were closed with butyl-rubber stoppers and sealed with aluminium crimps to avoid any possible gas exchange and oxygen presence. Cultures have been incubated in the dark at 42°C, and biogas and volatile fatty acids (VFA) productions have been measured to metabolically characterise the microorganisms. The isolates were kept active by continuous subcultures (approximately every 30 days), while long-term storage at −80°C was performed by adding 20% glycerol (v/v) suspension.

In order to obtain a protein-acclimated culture, inoculum from the F digester was diluted with BA medium at 20:80 v/v ratio, in 120 mL serum bottles with a working volume of 40 mL. Re-inocula from one generation to the following have been made every 21 days following the 20:80 v/v ratio in BA medium, using 8 g/L casein as carbon source and 0.2 g/L YE to sustain bacterial growth. To verify the activity of the culture, biogas production has been routinely measured along with the acclimatation strategies as described below.

### 2.5 Bioaugmentation experiment set up

The bioaugmentation experiment was applied on an AD system selectively fed with maize silage, casein, and their co-digestion. The inoculum used to start up the batches derived from a mesophilic lab-scale reactor in Thessaloniki (Greece), fed with bovine manure as the main substrate. The characteristics of the inoculum utilised for the experiment are described in **Table 1**.

**Table 1:** Characteristics of the inoculum and substrates (NA: not available; DM: dry matter)

| Parameters | Unit | Inoculum | Maize silage | Casein |
| --- | --- | --- | --- | --- |
| TS | (% of DM) | 6.52 ± 0.06 | 37.60 ± 0.31 | 91.59 ± 0.21 |
| VS | (% of DM) | 4.72 ± 0.01 | 36.71 ± 0.24 | 90.12 ± 0.19 |
| pH |  | 8.05 |  |  |
| VFA | (mg/L) | 138.79 ± 9.95 |  |  |
| NH<sub>4</sub><sup>+</sup>-N | (mg/L) | 4692.30 ± 100.98 | NA | NA |

Four sets of cultures were used for the bioaugmentation. A hydrogenotrophic archaea, *Methanoculleus bourgensis* MS2 (DSM3045), and a proteolytic bacterial strain, *Pseudomonas lundensis* DSM6252, were purchased from DSMZ GmbH (Leibniz Institute, Germany). *M. bourgensis* MS2 was grown at 37°C in 120 mL (40 mL working volume) anaerobic batch reactors in the medium DSMZ 332, with the addition of H_2_ and CO_2_ gas mixture (80/20, v/v) as substrate. Growth was weekly monitored by measuring methane production and optical density (OD_600nm_). *P. lundensis* DSM6252 was aerobically grown in NB at 25°C. A co-culture enriched with *Tissierella* species, obtained in the current work, was also selected for its proteolytic capabilities. At last, the third generation of the casein-acclimated culture obtained as discussed in section 2.4 was also tested.

Batch reactors with an organic loading (OL) of 4 g VS/L were set up in triplicate using 120 mL bottles, operating at inoculum to substrate (I:S) ratio of 9.5:1. Substrates were mixed with 24 g of mesophilic inoculum and an amount of distilled water required to reach the final working volume of 30 mL. Bioaugmentation cultures have been added to a concentration of 10% (v/v) of the final working volume. To establish anaerobic conditions, both liquid and headspace were flushed with N_2_ for 3 min each. The bottles were immediately closed with rubber stoppers, sealed with aluminium crimps, and incubated at 42 °C. Once per day the bottles were manually shaken to keep the solution well homogenised and every two days biogas production was monitored using the water displacement method. Biogas composition in terms of H_2_, CO_2_ and CH_4_ was measured by a gas chromatograph, and values expressed under the standard conditions of temperature and pressure, i.e. at a temperature of 0°C and a pressure of 1 atm. Benchmark experiments, containing only inoculum and distilled water, were also performed. Methane values were expressed in mL CH_4_/g VS.

### 2.6 Analytical methods and statistical analysis

Total solids (TS), volatile solids (VS), total Kjeldahl nitrogen (TKN) and pH were determined according to APHA Standard Methods for the Examination of Water and Wastewater (APHA, 2005).

CH_4_, H_2_ and CO_2_ content in biogas were determined three times per week using a gas chromatograph (490 Micro GC, Agilent Technologies, CA, USA) equipped with a thermal conductivity detector (TCD) and two different capillary columns, one using argon as carrier gas and the other using helium, operating at 145°C, 30 psi and 100°C, 28 psi, respectively. Data were analysed by SOPRANE 2 software (S.R.A. Instruments, France). For VFA analysis, samples were previously filtered using 0.22 μm cellulose acetate membrane and analysed through Shimadzu Nexera HPLC system, equipped with a RID-10A refractive index detector (Shimadzu, Kyoto, Japan). The chromatographic separations were performed using a Phenomenex Rezex ROA-Organic Acid H^+^ (8%) column (300 mm × 7.8 mm) set at 65°C. The analysis was performed at a flow rate of 0.6 mL/min using isocratic elution, with 5 mM H_2_SO_4_ as a mobile phase. VFA Mix 10mM (Sigma-Aldrich) was used as a reference standard. Analytes were identified by comparing their retention times and the concentrations were calculated by exploiting calibration curves of the corresponding external standard. All the determinations were performed in triplicate.

Statistical evaluation of the data set was also assessed using RStudio v3.6.3 (https://www.rstudio.com/), and the fmsb package v0.7.3 (https://cran.r-project.org/web/packages/fmsb/index.html). A *t* test assuming a normal distribution of data has been performed to evaluate statistical significance of methane yield of single conditions compared to the others. The *p*-value has been corrected considering non-independent tests, through FDR (false discovery rate) (Benjamini and Hochberg, 1995).

## 3. Results and discussion

### 3.1 Microbiome analysis of the digesters

A preliminary outline of the microbial composition of hydrolysis (H), fermentation (F) and post-fermentation (P) mesophilic digesters of a full-scale biogas plant located in Chiari (Brescia, Italy) was obtained by 16S rRNA gene sequence analysis. Total raw data includes 142566, 121276 and 102521 filtered reads for H, F and P, respectively. Up to 76% of the microbial community consists of Bacteria, with Firmicutes as the most abundant phylum (62-65%) followed by Bacteroidetes (18-27%) and Euryarchaeota (6-17%); furthermore, also Synergiststetes and Deinococcus-thermus are identified at low levels (2 and 1%, respectively) (**Supplementary Data**). Euryarchaeota is the unique phylum of the Archaea domain with a marked difference in abundance across the three microbial systems, with the highest value (17%) detected in the post-fermenter. This amount highly exceeds the one in the fermenter, confirming the continuous and intensive activity of methanogenic archaea also in the post-fermentation reactor.

The eighteen most abundant families are reported in **Figure 1a**. With the exception of Oscillospiraceae, which peaks at 25%, the most common families are Clostridiaceae, Methanomicrobiaceae, Erysipelotrichaceae, Dysgonomonadaceae and Marinilabiliaceae, with values ranging from 6 to 12% (**Figure 1a**). Since the taxonomy of the family is not strictly related to the metabolic function of the microorganism, a classification at the genus level is required, resulting more valid from a functional perspective. Despite differences in microbial communities of three reactors, a microbial core composed by *Ercella*, *Methanoculleus*, *Clostridium* and *Proteiniphilum* is shared by all digesters, representing half of the community in the post-fermenter (**Supplementary Data**). In fact, the hydrolytic and fermentative reactors are characterised by similar microbial compositions. Most of the OTUs did not have a high level of similarity with reference sequences in the public database. For this reason, only few OTUs can be tentatively classified to the species level, and the majority of them can be only assigned to the family level. Unidentified OTUs belonging to Marinilabiliaceae and Erysipelotrichaceae are predominant in H and F, followed by *Prevotella* sp. OTU15, *Clostridium* sp. OTU20, *Urmitella* sp. OTU17 and Bacteroidales sp. OTU22. Other OTUs are common and equally abundant in H, F and P such as Oscillospiraceae sp. OTU01 and OTU12, Erysipelotrichaceae sp. OTU03 and *Methanoculleus palmolei* OTU02. Considering the highly divergent functions of the different reactors, considerable variations in OTUs distribution were expected and can be observed between H and P digesters (**Figure 1b**). Moreover, P was also found to be rich in *M. bourgensis* OTU13 and Clostridiaceae and Bacteroidales species. *M. bourgensis* is affiliated to Methanomicrobiaceae family and it is frequently reported to play a central role in biomethanation (Granada et al., 2018). It is worth noting that the dominance of the *Methanoculleus* identified in these reactors indicates that the predominant methanogenic pathway was hydrogenotrophic. This is in agreement with studies of microbial population of biogas plants treating protein-rich substrates (Tian et al., 2019b; Treu et al., 2019) as the ammonia released by their digestion drives the methanogenic process from the most common acetoclastic pathway to the hydrogenotrophic one (Tian et al., 2018b).

**Figure 1:**
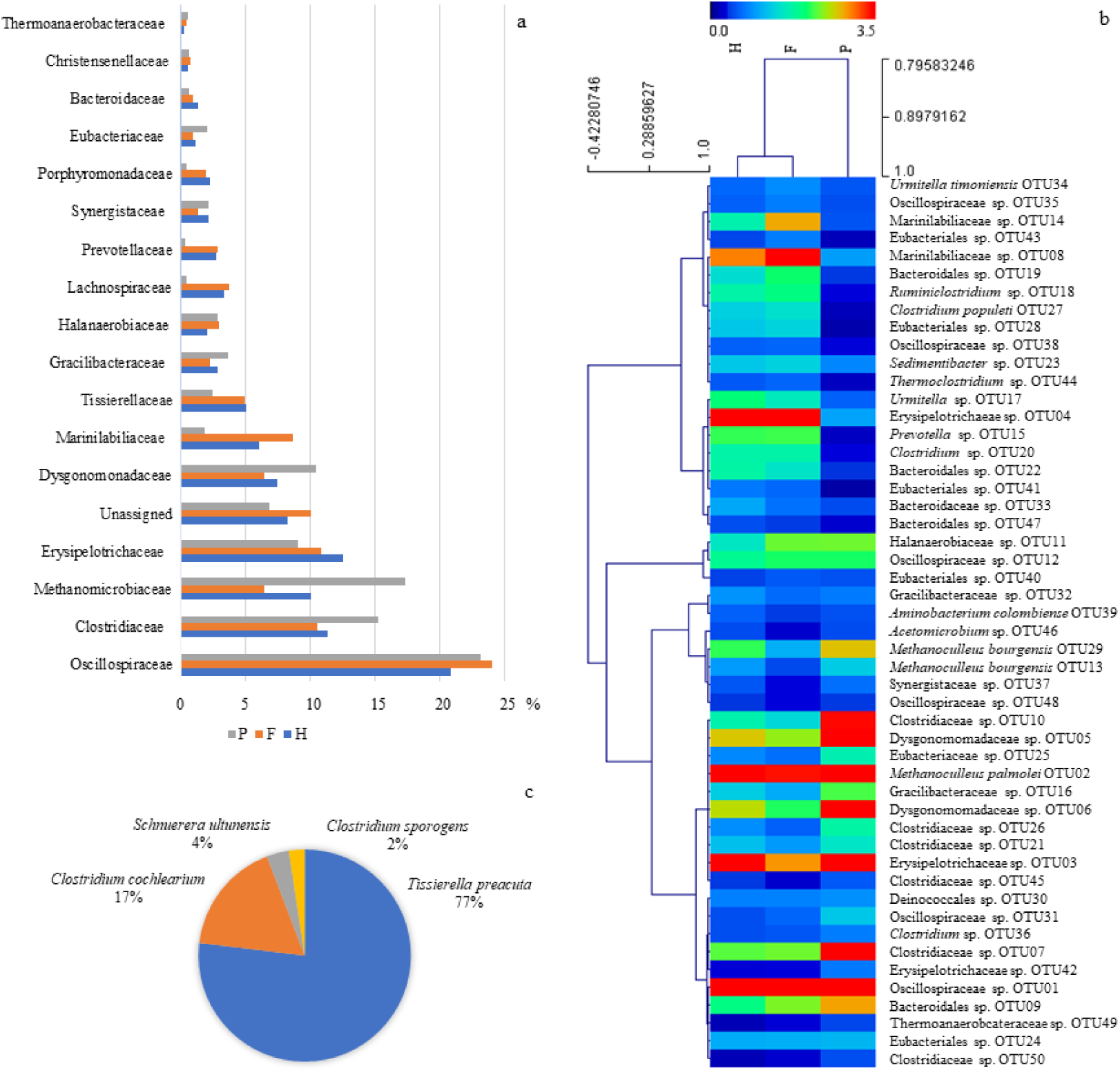
(**a**) Phylogenetic assignment based on 16S rRNA sequence analysis related to families (%) in H, F and P digesters. (**b**) Heatmap of OTUs abundance in each reactor and (c) species composition of the enriched co-culture used for bioaugmentation.

### 3.2 Isolation and characterisation of enriched microbial consortia and single species

It is ascertained that, towards a deeper knowledge of the anaerobic microbiome, culture-based techniques are needed to have a more detailed phenotypic characterisation of the species. The highest number of CFUs (nearly 3*10^6^ CFU/mL) was detected on NB and SM media, whereas lower CFU values (5*10^5^ CFU/mL) were obtained from acetic acid. Representative isolates from the three media were then selected for further studies and deeper characterisation.

A total of 40 isolates were obtained in single culture or co-culture and tested for their ability to produce VFA, H_2_, CO_2_, and CH_4_ from different carbon sources. Isolates having the phenotypic properties of interest were selected and identified by 16S rRNA sequencing (**Table 2**). All of them were assigned to Firmicutes phylum, and, more specifically, to classes of Clostridia (four isolates), Bacilli (two isolates) and Tissierellia (three isolates). All the selected isolates were able to produce various levels of CO_2_ and H_2_ from different substrates. H_2_ production varies from 1.8 to 15% whereas CO_2_ from 5.7 to 25.7% (**Table 2**). The highest levels of H_2_ were produced by the isolates L6 and L21, as well as by the co-culture I4. Using acetate as the carbon source, *Schnuerera ultunensis* L21 produced 25.7% of CO_2_ and 12.2% of H_2_. This agrees with other papers describing *S. ultunensis* as able to convert acetate into CO_2_ and H_2_ through the Wood-Ljungdahl pathway (Manzoor et al., 2016; Schnurer et al., 1996). This microbe is indeed a SAOB, as it establishes a syntrophy with hydrogen-consuming archaea (Hattori et al., 2008). It is also commonly abundant in anaerobic digestion systems releasing ammonia from substrates degradation, supporting the methane production via hydrogenotrophic pathway, usually dominated by *Methanoculleus* spp. (Tian et al., 2018b). Significant quantities of CO_2_ and H_2_ were also released by *Caproiciproducens galactitolivorans* L6, supporting what previously described by Kim et al., 2015, according to which this species can convert skim milk into simple molecules, such as gases and VFA. *Bacillus thermoamylovorans* I6 mostly produced CO_2_ and limited amounts of H_2_, in agreement with other works reporting *B. thermoamylovorans* to boost H_2_ production from biomass in co-cultures with *Clostridium* sp. strains (Cabrol et al., 2017).

**Table 2.**
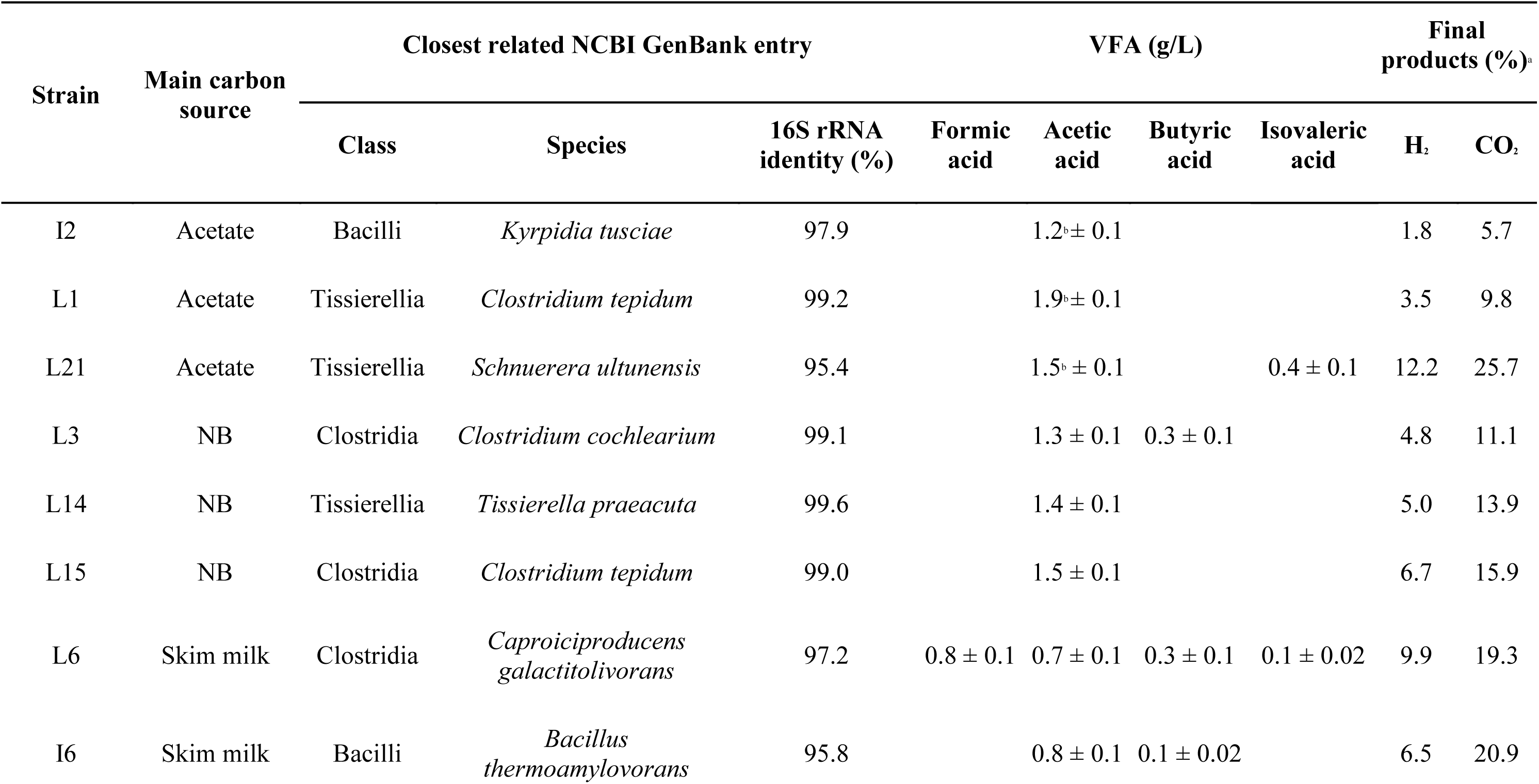
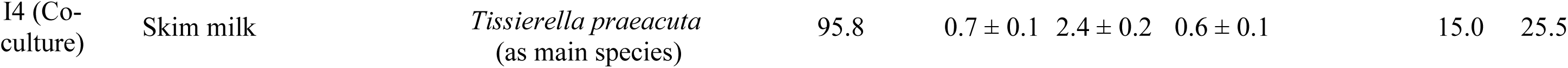
16S rRNA sequencing of selected bacterial strains anaerobically isolated from acetate, SM and NB and their technological properties in terms of VFA and gas production from different carbon sources. ^a^ Partial pressure. ^b^Residual acetic acid after bacterial growth on acetate.

Despite the strong variability observed, all the selected strains produced a significant amount of VFA (**Table 2**), with the exception of *Kyrpidia tusciae* I2. Regarding the VFA profile, acetate was the main product, but also butyric, formic and isovaleric acids were identified in some cultures. *C. galactitolivorans* L6 produced the largest VFA profiles as formic, acetic, butyric and isovaleric acids were detected in the spent broth containing skim milk as the main carbon source.

1. *C. cochlearium* L3, a potential proteolytic organism involved in cheese spoilage (Gupta et al., 2020), produced butyric and acetic acid from NB. This is in accordance with other papers describing this species as producer of both VFA from various protein-rich carbon sources (Nzeteu et al., 2018), which are abundantly present in the NB formulation. As expected, isovaleric and acetic acids were produced by *S. ultunensis* L21 in agreement with other studies (Schnurer et al., 1996).

A co-culture named I4 showed promising technological traits, being able to produce the highest levels of H_2_ and CO_2_ (15 and 25.5%, respectively) from skim milk (**Table 2**). Furthermore, the co-culture produced interesting amounts of formic, acetic and butyric acids. Moreover, its proteolytic activities were confirmed on plate (data not shown), suggesting the potential of these microbes to be adopted as co-culture for anaerobic digestion of protein-rich organic waste streams. High throughput 16S rRNA amplicon sequencing of this co-culture disclosed that it is mostly composed by *Tissierella praeacuta* (**Figure 1c**). The total number of 16S rRNA amplicon reads for the sample of interest was 264910, with an average length of 289 bp. In the selected culture, the OTU assigned to *T. praeacuta* (100% identity) showed a 66.9% relative abundance, whereas C. *cochlearium* (100% identity) and S. *ultunensis* (94.9% identity) were also assigned to OTUs with 14.6% and 2.8% relative abundance, respectively.

### 3.3 Bioaugmentation strategy to optimise AD of protein-rich substrates

A previous experiment of co-digestion of maize silage and casein involving the same inoculum used in this work and the addition of the proteolytic bacteria *P. lundensis* DSM6252 resulted in a methane yield decrease (data not shown). This finding points out that bioaugmentation with a proteolytic species does not favour the biogas production, rather causes the release of ammonia provoking the inhibition of the AD process. However, since hydrogenotrophic methanogens, specifically *Methanoculleus* spp. (Tian et al., 2018a), have been reported to be the most resistant archaea to ammonia, in the present work a bioaugmentation with a proteolytic bacterium in combination with *M. bourgensis* MS2 has been assessed. *P. ludendsis* DSM6252 as well as the co-culture I4 were tested to enhance protein hydrolysis, while the ammonia-tolerant methanogen, or a community acclimated to a double casein OL, was adopted to increase the methanation rate.

Casein and maize silage were here selected as representative substrates for proteinaceous and agricultural AD feedstocks, respectively. The several bioaugmentation approaches adopted are briefly described below. The choice of *P. lundensis* DSM6252 leans on previous findings about its relevance in the degradation of protein substrates, mainly derived from cheese whey and wastes (Fontana et al., 2018). Furthermore, the functional analysis of a MAG (Metagenome-Assembled Genome) taxonomically assigned to *P. lundensis* DSM6252 reported the presence of a complete β-oxidation pathway and 21 proteases (Fontana et al., 2018; Morales et al., 2005). Likewise, since the prevalent species in the co-culture I4 (**Figure 1c**), *T. praeacuta*, was predicted to be proteolytic, this culture was used to compare its effectiveness with that of the collection strain of *P. lundensis* DSM6252. Moreover, the presence of *C. cochlearium* in the same culture (**Figure 1c**) must be taken into account, given its implication in degradation of dairy products. As a parallel strategy, the bioaugmentation with the culture adapted to casein was meant to potentially improve the AD performance, since a selected casein-degrading consortium is expected to have better performances when casein is the sole carbon source. The choice of *M. bourgensis* MS2 lies in its ability to thrive under high ammonia concentrations without hampering the methane yield, demonstrating high resilience in conditions that can be stressful for several methanogens (Tian et al., 2018a; Yan et al., 2020).

#### 3.3.1 Batch reactors AD of casein under different bioaugmentation approaches

The four bioaugmentation strategies above described, namely M (*M. bourgensis*), PM (*P. lundensis* and *M. bourgensis*), EM (enriched co-culture I4 and *M. bourgensis*), and A (casein acclimatised culture), were compared to the benchmark not bioaugmented (NB) in batch reactors for the production of biogas from maize silage, casein and their combination. Their methane productivity was monitored in a time-course experiment (**Figure 2**), together with pH and VFA profiles, as well as TKN levels (**Figure 3**).

**Figure 2:**
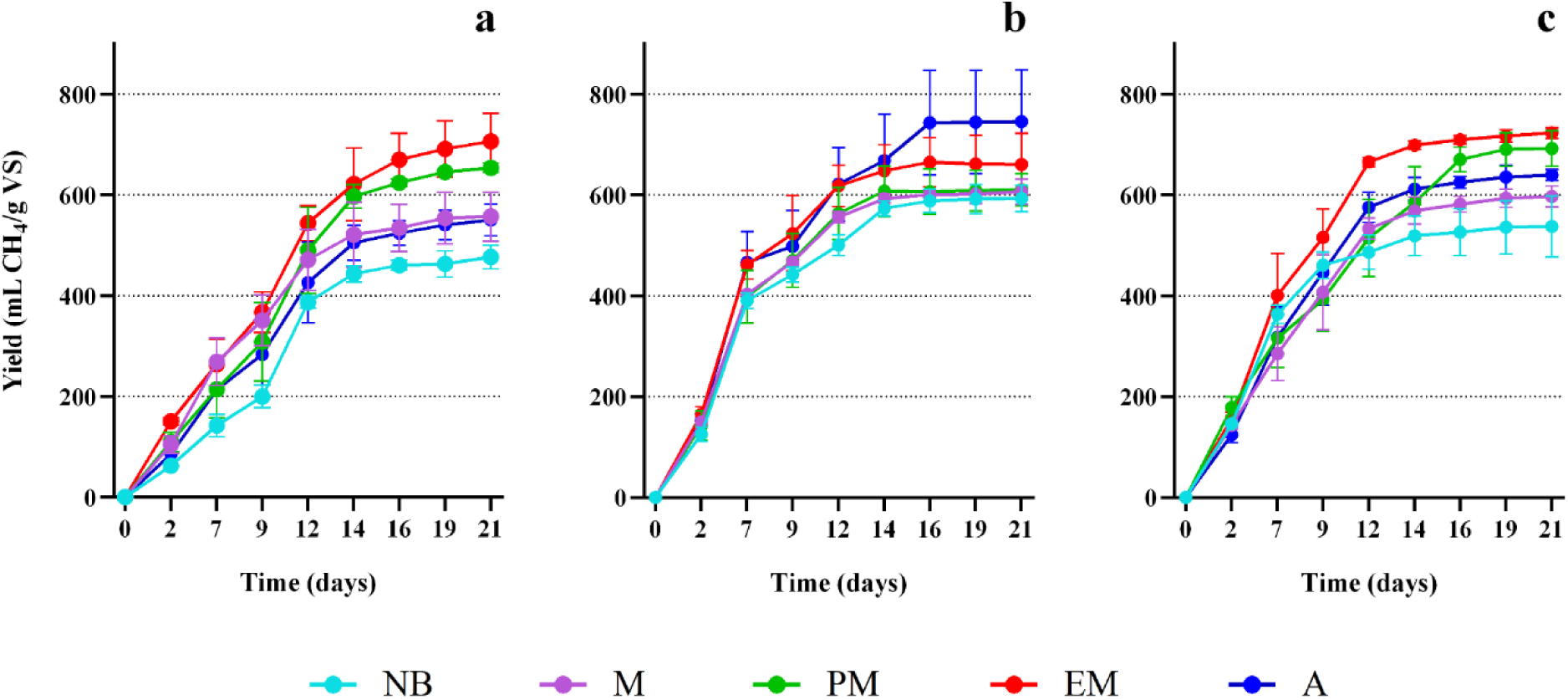
Methane production yields (mL CH_4_/g VS) of batch reactors fed with maize silage (**a**), casein (**b**) and their combination (**c**). The different conditions are reported as: not bioaugmented (NB), *M. bourgensis* (M), *P. lundensis* and *M. bourgensis* (PM), enriched culture and *M. bourgensis* (EM), casein acclimatised culture (A). Error bars denote standard deviation from the mean of triplicate measurements (*n*=3).

**Figure 3:**
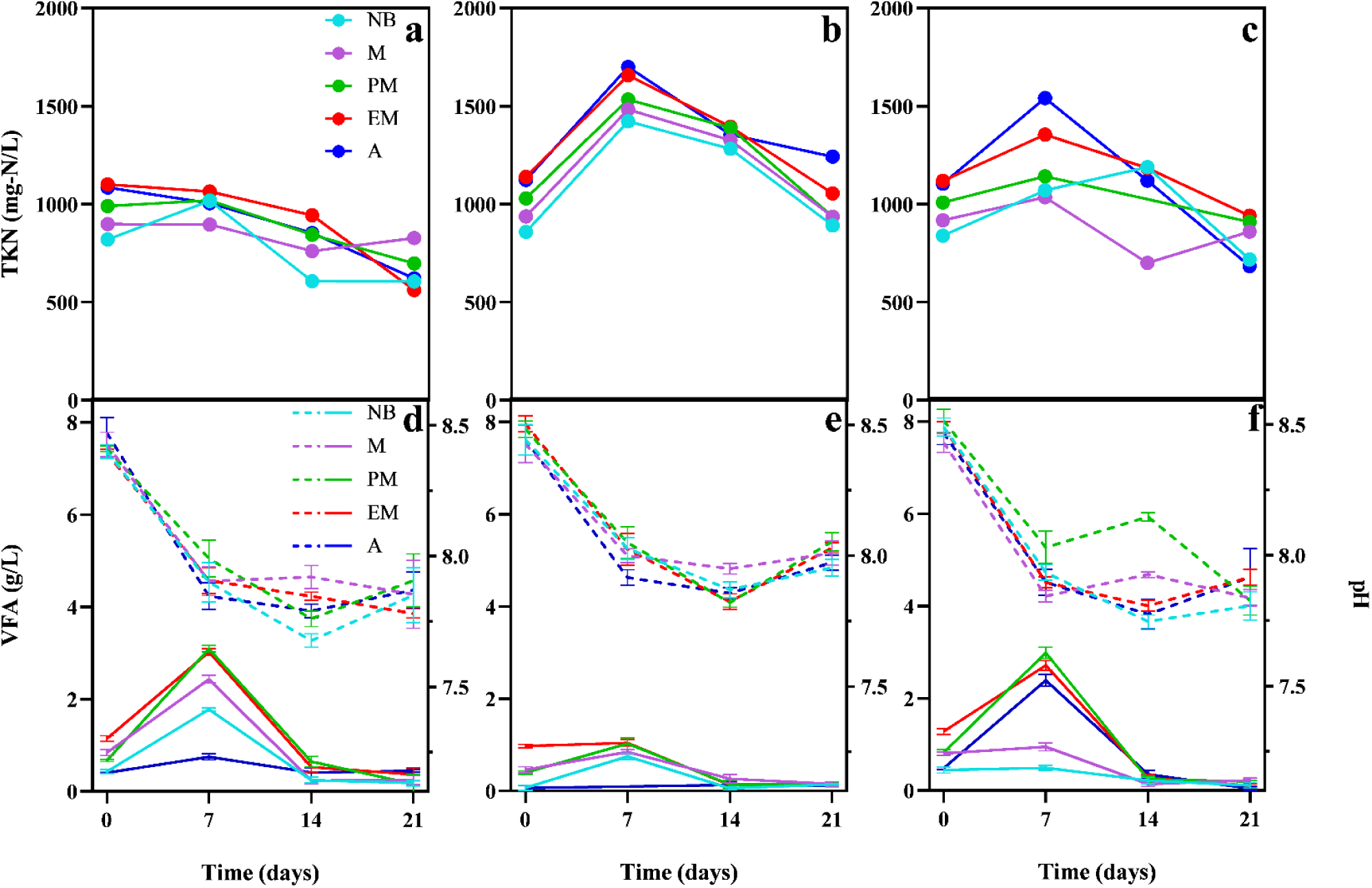
TKN (mg-N/L), total VFA (full lines) and pH (dashed lines) values during AD of maize silage (**a**, **d**), casein (**b**, **e**) and their combination (**c**, **f**).

Intermediate products such as VFA and the process parameter pH were measured once per week. The pH of the reactors fluctuated from 8.5 to 7.65 with a decreasing trend (**Figure 3d-f**), hence remaining in the optimal range for methane production (Kougias and Angelidaki, 2018). All the experimental conditions have similar trends and evidence a reduction in pH as a consequence of substrate degradation and VFA production. Specifically, it is noteworthy that a pH reduction to approximately 7.7 corresponds to an increase in methane production till it reaches the steady state (days 14-16) (**Figure 2**), remaining in the optimal pH for an efficient methanogenesis of around 6.5–8.2 (Kougias and Angelidaki, 2018). In batch settings, VFA generally accumulate in the first phase of the process and sharply decrease in the following steps when methanogenesis occurs very rapidly. This is in agreement with the total VFA profile detected over time also in this study. The VFA profile is dominated by acetic acid, followed by propionic and valeric acids (**Supplementary Data**).

Although the initial kinetics of all the experimental conditions per each substrate appear to be similar, differences in methane yield become more evident while reaching the stationary phase (**Figure 2**). In all three tested cases, bioaugmentation strategies were revealed to effectively increase biogas production.

The methane yield of the NB condition from maize silage (**Figure 2a**) is 477±44 mL CH_4_/g VS, lower than those obtained in the bioaugmented reactors. Particularly, the highest methane production with this substrate was achieved in the case of the EM bioaugmentation, which peaked at 707±55 mL CH_4_/g VS, corresponding to 1.48-fold the NB control condition. EM is followed by PM, which induced a production 37% higher than NB (654±9 mL CH_4_/g VS). Overall, both conditions resulted in methane yields statistically higher than those detected for NB, M and A. In fact, both involve the bioaugmentation with a combination of *M. bourgensis* and a proteolytic bacterium/consortium: the effect of the high release of ammonia from the proteolytic activity can be mitigated by the resilience of *M. bourgensis*, which in turn drives the archaeal community towards a more efficient methanogenesis (Tian et al., 2019a).

Reactors fed with maize silage evidenced a similar trend for M and A, with a methane yield increase of 17 and 15%, respectively, when compared to the NB condition (**Figure 2a**). Considering the high specialisation of the culture A for a different, protein-rich feedstock (casein), the main contribution to the system was due to methanogenic activity, which appears to be comparable to that measured in the case of M bioaugmentation. This finding is confirmed since those differences in methane productions detected in the case of NB, M or A are not statistically significant. The greater bioaugmentation effects observed with EM and PM cultures are likely due to their microbial architecture, since both contain hydrolytic and proteolytic bacteria which can efficiently degrade macronutrient added to the reactors. Focusing on EM, results may indicate a higher degradation rate of maize silage, resulting in larger nutrient availability, and consequently in a greater methane yield. Even though methanogenesis is the rate-limiting step, it is possible that the high presence of *M. bourgensis* in the bioaugmentation culture pushed toward the higher consumption of molecules generated in the other AD steps by bacterial strains. VFA and pH profiles seem to support this finding since the highest values of total VFA were detected in the bioaugmented batches, and pH levels decreased accordingly (**Figure 3d**). Nitrogen content did not change significantly in the different inoculation strategies (**Figure 3a**).

Methane production from casein was faster than in maize silage with, on average, almost 66% of the biogas that was already produced after 7 days of incubation (**Figure 2b**). As expected, the NB condition resulted in methane yield higher than that obtained from maize silage (p < 0.05). Moreover, the most efficient bioaugmentation was found to be with the A culture (745±103 mL CH_4_/g VS, +26% on average). Significantly lower methane yield and productivity values were displayed by the EM approach (661±62 mL CH_4_/g VS, +11%). The proficient proteolytic activities of both A and EM were evident considering the increase in total nitrogen content (**Figure 3b**) and the highest total VFA profiles (**Figure 3e**). Noteworthy, both bioaugmentations were more productive than the batch experiments inoculated with PM and M. This suggests a lower hydrolytic activity of *P. lundensis* (PM) at such high OL of casein, in comparison to the one of the enriched co-culture I4 (EM).

Checking biochemical parameters, higher TKN values were expected, since casein contains much more nitrogen than maize silage. Total VFA are similar in all the conditions, except for the reactors bioaugmented with A (**Figure 3e**), which shows lower values. This further indicates that the VFA produced during the acidogenesis phase were immediately converted into CH_4_, especially when A was used for bioaugmentation, confirming its higher casein conversion efficiency.

Considering now the co-digestion of casein and maize silage, EM and PM cultures once again triggered biogas productions higher (p < 0.05) than those of the not bioaugmented reactors, with increments in methane yields corresponding to 34 and 19%, respectively. As such, the presence of *P. lundensis* or the newly isolated enriched culture was efficient in supporting the feedstock hydrolysis (**Figure 2c**) and then the production of methane in a co-digestion context.

As expected, the bioaugmentation with A, which showed the highest potential in CH_4_ production from casein, produced lower values once in presence of both maize silage and casein. The methane yield (640±11 mL CH_4_/g VS) was indeed approximately intermediate between those observed with maize silage (550±32 mL CH_4_/g VS) and with casein (745±103 mL CH_4_/g VS) alone. The same observation is also found in the TKN trend (**Figure 3c**).

On the contrary, the bioaugmentation with M did not result in increased methane yields compared to the benchmark. Overall, it is noteworthy that the use of a mixed culture with an archaeon and a hydrolytic bacteria avoided the inhibition of the system. In fact, their methane yields never fell behind the NB ones. Similarly, TKN has intermediate values between the single feedstocks digestion (**Figure 3a,b**). Moreover, the steepest decrease in TKN measured in the A condition after 7 days may indicate a certain incorporation of the ammonia in the cell biomass, thus suggesting a higher growth rate of the community. Considering that the co-digestion of maize silage and casein can be representative of many agricultural anaerobic digesters treating different feedstocks, the results obtained in this condition are the most interesting ones and indicated that the EM and PM bioaugmentations were the most promising. In fact, they were effective in alleviating the stressing conditions triggered by casein supplementation in the benchmark AD (NB).

## 4. Conclusions

In this study, newly isolated strains and a co-culture from an anaerobic digester were investigated for their potential in biogas production. A combination of genomic and biochemical analysis was used to characterise the selected inoculum and co-cultures. The bioaugmentation approach was successful in boosting methane production from difficult substrates. The best results were obtained from the casein consortium and proteolytic co-culture with *M. bourgensis* for the digestion of casein and its co-digestion with maize silage. Such bacterial and archeal combination is currently under study on continuous reactors to further assess the feasibility of this bioaugmentation strategy at larger scale towards an efficient and stable AD of protein-rich feedstocks.

Supplementary data related to this work can be found in the online version of the paper.

## Acknowledgments

We acknowledge Valentino Pizzocchero (MSc) for technical assistance in GC and HPLC analysis. Fundings: This project was supported by Fondazione Cariverona under the project “Più-BIOGAS APP”, the funding BIRD210708/21, the project ‘‘Sviluppo Catalisi dell’Innovazione nelle Biotecnologie” (MIUR ex D.M.738 dd 08/08/19) of the Consorzio Interuniversitario per le Biotecnologie” (CIB) and by the project LIFE20 CCM/GR/001642 – LIFE CO2toCH4 of the European Union LIFE+ program. The microGC (Agilent Technologies, CA, USA) was funded by Progetto di Eccellenza “Centro per l’Agricoltura, la Sostenibilità e gli Alimenti” (CASA), CUP C26C18000190001, MIUR, Italy.

## CRediT author statement

**Sara Agostini**: Investigation, Data Curation, Writing - Original Draft, Visualisation. **Luca Bucci**: Investigation, Data Curation, Writing - Original Draft, Visualisation. **Davide Doni**: Resources. **Paola Costantini**: Resources. **Ameya Gupte**: Investigation. **Bettina Müller**: Resources. **Fabrizio Sibilla:** Resources. **Marina Basaglia**: Resources. **Sergio Casella**: Resources. **Panagiotis G. Kougias**: Conceptualisation, Resources. **Stefano Campanaro**: Conceptualisation, Supervision, Resources, Writing - Review & Editing, Funding acquisition. **Lorenzo Favaro**: Conceptualisation, Validation, Supervision, Resources, Writing - Review & Editing, Funding acquisition. **Laura Treu**: Conceptualisation, Methodology, Supervision, Resources, Writing - Review & Editing, Funding acquisition.

## Notes

### Competing Interest Statement

The authors have declared no competing interest.

